# Characteristic Localization of Neuronatin in Rat Tissues

**DOI:** 10.1101/447169

**Authors:** Naoko Kanno, Saishu Yoshida, Takako Kato, Yukio Kato

**Author notes:** Corresponding Author: Yukio Kato, Meiji University, 1-1-1 Higashi-Mita, Tama-Ku, Kawasaki, 214-8571 Kanagawa, Japan.

## Abstract

Neuronatin (*Nnat*) is expressed in the pituitary, pancreas, and other tissues; however, the function of NNAT is still unclear. Recent studies have demonstrated that NNAT is localized in the sex determining region Y-box 2-positive stem/progenitor cells in the developing rat pituitary primordium and is downregulated during differentiation into mature hormone-producing cells. Moreover, NNAT is widely localized in subcellular organelles, excluding the Golgi. Here, we further evaluated NNAT expression and intracellular localization in embryonic and postnatal rat tissues such as the pancreas, tongue, whisker hair follicle, and testis. Immunohistochemistry showed that NNAT was localized in undifferentiated cells (i.e., epithelial basal cells and basement cells in the papillae of the tongue and round and elongated spermatids of the testis) as well as in differentiated cells (insulin-positive cells and exocrine cells of the pancreas, taste receptor cells of the fungiform papilla, the inner root sheath of whisker hair follicles, and spermatozoa). Additionally, NNAT showed novel intracellular localization in acrosomes in the spermatozoa. Because the endoplasmic reticulum (ER) is excluded from spermatozoa and sarco/ER Ca^2+^-ATPase isoform 2 (SERCA2) is absent from the inner root sheath, these findings suggested that NNAT localization in the ER and its interaction with SERCA2 were cell-or tissue-specific properties.

## Introduction

Neuronatin (NNAT) is a small protein consisting of two splice variants encoding 81 (NNATα) and 58 amino acids (NNATβ) and is well conserved evolutionally.^1,2^ NNAT has attracted attention from many researchers because of its high expression in the neonatal brain^3-5^ and pituitary gland.^6^ However, the role of NNAT *in vivo* is poorly understood. Recently, we found that NNAT first appears in and occupies stem/progenitor cells in the rat embryonic pituitary primordium and is then downregulated during differentiation into mature hormone-producing cells, suggesting that NNAT may be involved in pituitary development^7^ Moreover, we found that NNAT localizes in various subcellular organelles in addition to the endoplasmic reticulum (ER), as reported previously.^8^

Several important roles of NNAT have been reported in the past decade. For example, Lin et al. demonstrated that NNAT promotes neural differentiation of embryonic stem cell and that NNAT interacts with sarco/ER Ca^2+^-ATPase isoform 2 (SERCA2).^9^ Involvement of NNAT in the differentiation of keratinocytes^10^ and adipose tissue^11^ has also been suggested. Moreover, several studies have demonstrated diverse roles of NNAT,81213 such as insulin secretion, synaptic plasticity, calcium-induced cell migration, stress responses,^14,15^ and others.^16-18^ Thus, NNAT performs diverse functions according to its expression in particular tissues, although the details have not yet been reported. Furthermore, the specific expression of NNAT in various cell types within tissues has not been elucidated.

Accordingly, in this study, we analyzed the localization of NNAT-positive cells using immunohistochemistry in embryonic and postnatal tissues, with a particular focus on the tongue papillae, whisker hair follicles, testes, and pancreas. We re-evaluated the intracellular localization of NNAT in specific subcellular organelles other than the ER and the colocalization of NNAT with SERCA2. Our observations provide important insights into the function of NNAT in tissue development.

## Materials and Methods

### Animals

Male Wistar rats were housed individually in a temperature-controlled room under a 12-hr light/12-hr dark cycle. Determination of pregnancy was made by the observation of a vaginal plug on day 0.5 of gestation. Rats were euthanized by cervical dislocation under anesthesia. The present study was approved by the Institutional Animal Care and Use Committee of Meiji University and was performed based on NIH Guidelines for the Care and Use of Laboratory Animals.

### Immunohistochemical Analyses

Postnatal rats were administered anesthetic and then subjected to perfusion fixation using 4% paraformaldehyde in 20 mM HEPES. Tissues were appropriately dissected and placed in 4% paraformaldehyde in 20 mM HEPES overnight at 4°C, followed by immersion in 30% trehalose in 20 mM HEPES for cryoprotection. Tissues were then embedded in Tissue-Tek OCT compound (Sakura Finetek Japan, Tokyo, Japan) and frozen immediately. Frozen sections (4 or 10 μm thick) on glass slides (Matsunami, Osaka, Japan) were activated as necessary using Immunosaver (Nisshin EM, Tokyo, Japan) for 60 min at 80°C and washed three times in 20 mM HEPES (pH 7.5) containing 100 mM NaCl (HEPES buffer) for 15 min. This was followed by blocking with 0.4% Triton X-100 (Sigma-Aldrich Corp., Tokyo, Japan) and 10% fetal bovine serum in HEPES buffer (pH 7.5) for 60 min at room temperature. Sections were then incubated for 16 hr at 4°C with the following primary antibodies: rabbit IgG against mouse NNAT (1:250 dilution; ab27266; Abcam, Cambridge, UK), guinea pig IgG against human insulin (1:100 dilution; ab7842; Abcam), goat IgG against human sex determining region Y-box 2 (SOX2; 1:500 dilution; GT15098; Neuromics, Edina, MN, USA), mouse IgG against human E-cadherin (1:200 dilution; BD Bioscience, San Jose, CA, USA), and mouse monoclonal antibody against rat protein disulfide isomerase (PDI; 1:1000 dilution; ab2792; Abcam). After being washed with HEPES buffer three times, sections were incubated for 2 hr at room temperature with Cy3, Cy5, or fluorescein isothiocyanate (FITC)-conjugated donkey anti-rabbit, anti-guinea pig, or anti-goat IgG as secondary antibodies (Jackson ImmunoResearch, West Grove, PA, USA). This was followed by enclosing in Vectashield mounting medium containing 4’,6-diamidino-2-phenylindole (DAPI; Vector, Burlingame, CA, USA) along with observation using fluorescence microscopy BZ-9000 (Keyence, Osaka, Japan) and fluorescence confocal microscopy FV1000 (Olympus, Tokyo, Japan).

### Hematoxylin and Eosin (HE) Staining

Before staining, frozen sections (10 μm thick) on glass slides were washed with water for 15 min, stained for 10 min with hematoxylin (Wako, Osaka, Japan), washed with water for 1 hr, and stained with eosin (diluted 5-fold with 95% ethanol; Muto Pure Chemicals Co., Ltd., Tokyo, Japan). Finally, after dehydration, the sections were sealed for observation.

## Results

### Immunohistochemistry for NNATin the Pancreas

Previous studies have reported the expression of NNAT in the pancreas, an endodermal appendage;^8,19^ however, the relationship of NNAT with the development of the embryonic pancreas has not yet been examined. Hence, we investigated whether NNAT is present in the embryonic pancreas and reconfirmed the intracellular localization of NNAT in the adult pancreas.

HE staining of embryonic and adult pancreatic tissues was performed to show developing pancreatic islets and exocrine cells in addition to the pancreatic ducts and blood vessels (Fig. 1A–C). Immunohistochemistry for NNAT showed a small number of immunoreactive signals in the region of exocrine cells on embryonic day 16.5 (E16.5; Fig. 1A, D, E, white arrowhead), and granular signals were observed in pancreatic islets (Fig. 1B: PI; Fig. 1F, G: white arrowhead) and exocrine cells on E21.5 (Fig. 1B: EC; Fig. 1F, H: white arrowhead). The signal intensity was increased on postnatal day 60 (P60; Fig. 1C, I–K, white arrowheads). Strong tube-shaped signals were observed on P60 (Fig. 1I, yellow arrow). Double immunostaining for NNAT and E-cadherin, which is known to be expressed in the pancreatic duct, showed that cells with tube-shaped NNAT signals were negative for E-cadherin, except for E-cadherin-positive pancreatic ducts (Fig. 1l, M1–M3, yellow arrows and yellow arrowheads, respectively). Because NNAT has been reported to be expressed in the endothelial cells of the pancreatic aorta,^20^ these tube-shaped signals may represent pancreatic endothelial cells of the artery.

**Fig. 1.**
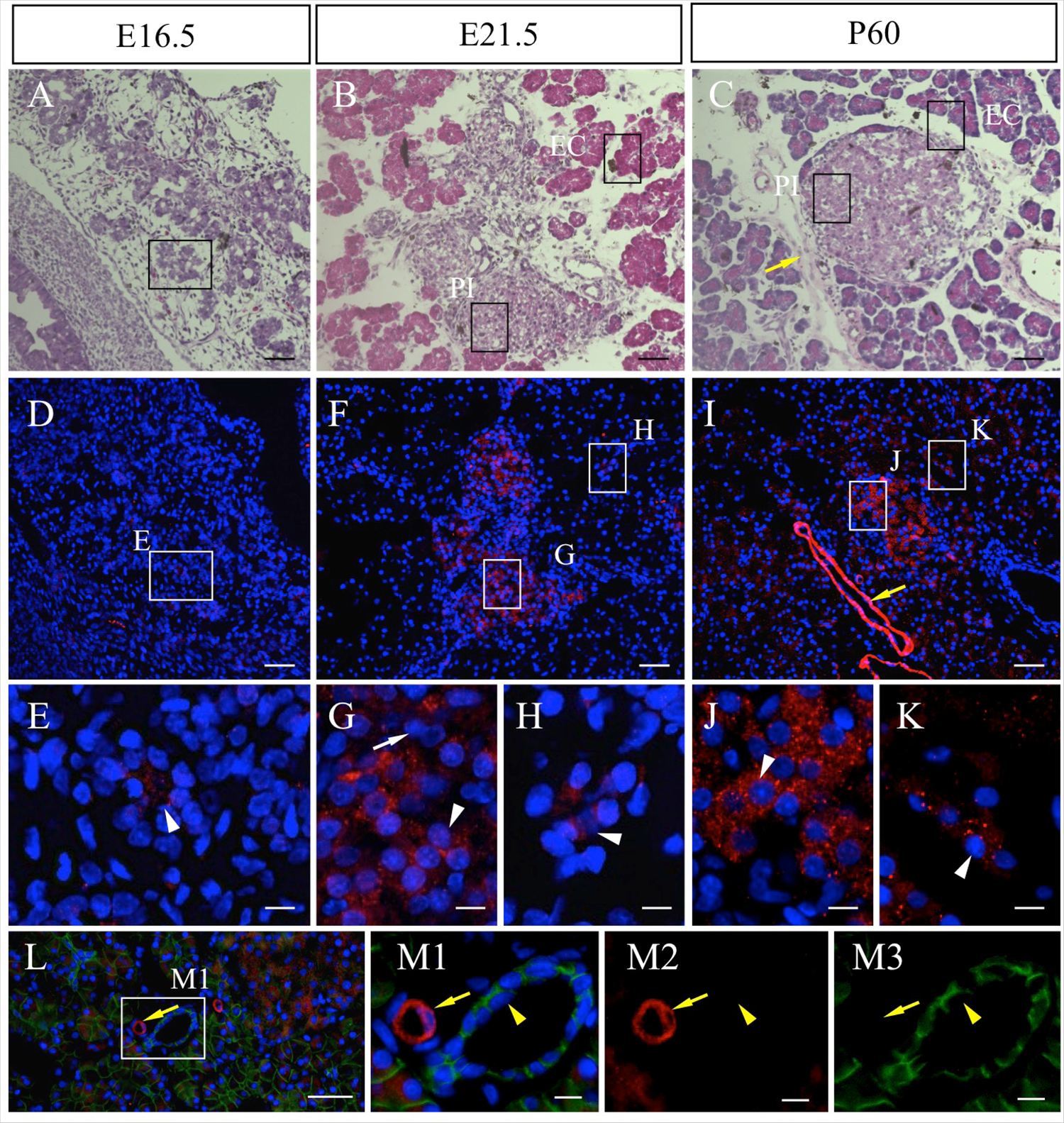
Localization of NNAT in embryonic and adult pancreatic tissues. Hematoxylin and eosin staining (A–C) and immunostaining for NNAT (D–K, red, Cy3) was performed using serial frozen sections prepared from embryonic tissues on embryonic day 16.5 (E16.5) (A, D) and E21.5 (B, F) and from adult tissues on postnatal day 60 (P60) (C, I). Boxes in A–C indicate areas adjacent to boxes in D, F, and i, respectively. Nuclei are stained with 4’, 6-diamidino-2-phenylindole (DAPI, blue). Double immunostaining for NNAT (red, Cy3) and E-cadherin (green, FITC) was performed on P60 (L, Ml–M3). Boxed areas in D, F, I, and I are enlarged in E, G, H, J, K, and M1–M3. White arrowhead: NNAT-positive cell, white arrow: NNAT-negative cell, yellow arrow: blood vessel, yellow arrowhead: pancreatic duct. Scale bars = 50 μm (A–C, D, F, I, L), 10 μm (E, G, H, J, K, M1–M3).

Next, we performed double immunostaining for NNAT and insulin because some NNAT-negative cells were detected in the pancreatic islets on E21.5 (Fig. 1G, white arrow). The results showed that insulin-positive cells had already emerged in the embryonic pancreas on E16.5 (Fig. 2A), but were negative for NNAT (Fig. 2B1–B4, arrows). In the pancreatic islets on E21.5 (Fig. 2C: PI), cells double positive for NNAT and insulin were observed, as were NNAT-single-positive and insulin-single-positive cells (Fig. 2D1–D4, open arrowheads, white arrowheads, and arrows, respectively). NNAT-positive cells were also detected in exocrine cells (Fig. 2C: EC, Fig. 2 D1–D4: white arrowheads on the lower side). In adult islets, cells positive for NNAT or insulin alone were still detected (Fig. 2E, F1-F4, white arrowheads and arrows, respectively), whereas most insulin-positive cells were positive for NNAT (Fig. 2E, F1–F4, open arrowheads). NNAT-positive cells were also observed in exocrine cells. Insulin-positive cells accounted for about 59% of pancreatic islet cells (Fig. 2G) and were mostly positive for NNAT on P60. Insulin/NNAT-double-positive cells accounted for approximately 96% of insulin-positive cells and 97% of NNAT-positive cells (Fig. 2G), showing a high proportion of colocalization.

**Fig. 2.**
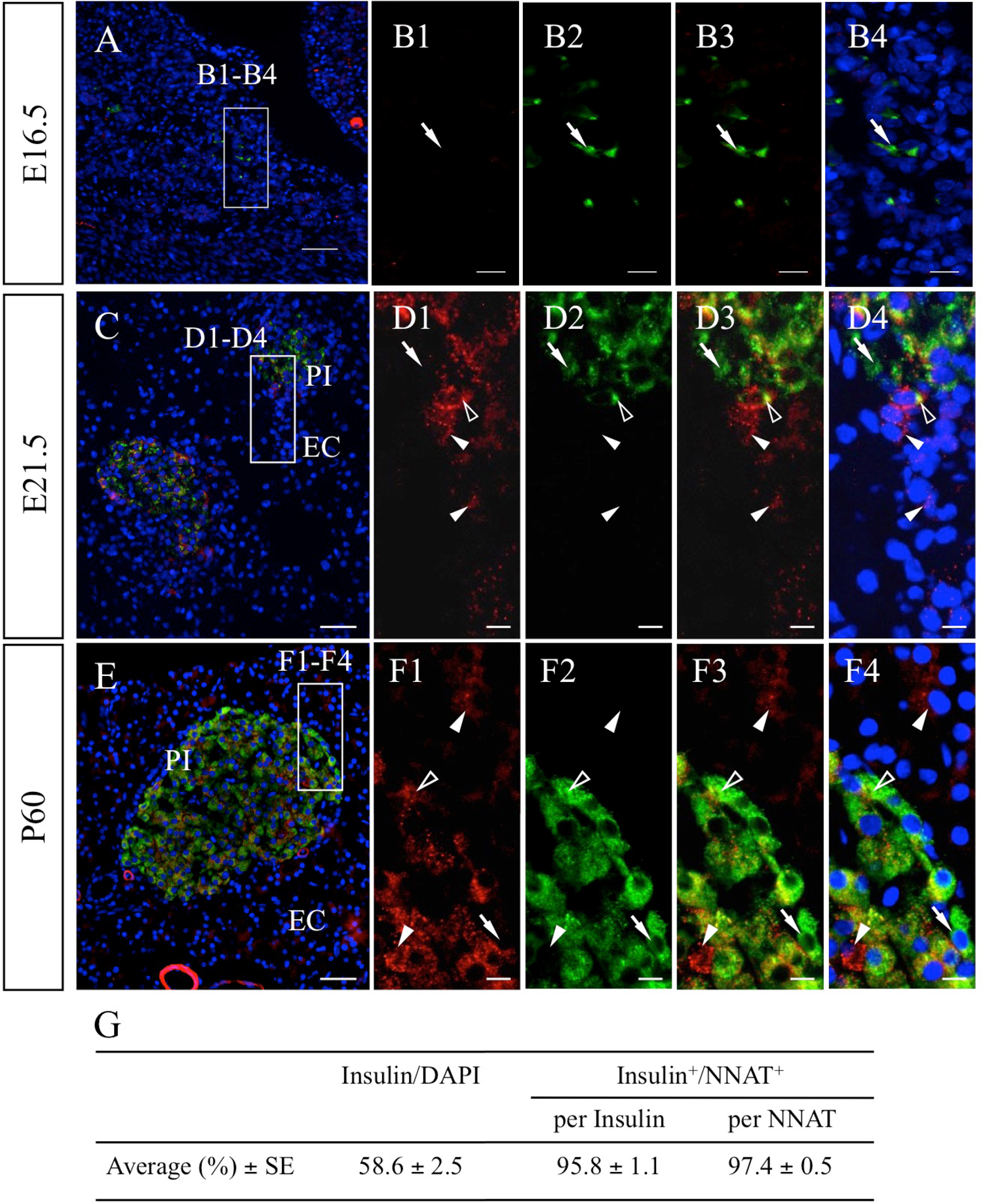
Immunohistochemistry for NNAT and insulin in embryonic and adult pancreatic tissues. Immunostaining for NNAT (red, Cy3) and insulin (green, FITC) was performed for frozen sections prepared from tissues on E16.5 (A), E21.5 (C), and P60 (E). Boxed areas in A, C, and E are enlarged in B1–B4, D1–D4, and F1–F4. The proportions of insulin-positive cells/DAPI and insulin/NNAT-double positive cells in insulin-positive and NNAT-positive cell populations were counted for six areas (129–495 cells each) and are listed in g. Data are presented as means ± SEs. White arrowhead: NNAT-single-positive cell, open arrowhead: NNAT/insulin-double positive cell, arrow: insulin-single-positive cell. PI: pancreatic islet, EC: exocrine cell. Scale bars = 50 μm (A, C, E), 10 μm (B1–B4, D1–D4, F1–F4).

In the pancreas, NNAT localizes in the ER of β-cells,^8^ However, in a previous study of the pituitary gland, we observed, for the first time, that NNAT was widely localized in subcellular organelles in addition to the ER.^7^ Consequently, we have re-examined subcellular localization of NNAT in the adult pancreas using an antibody against PDI, a marker of the ER (Fig. 3A). The results showed that, whereas most NNAT-positive cells in pancreatic islets were positive for PDI, some signals with positivity for NNAT only were also observed (Fig. 3B1–B5, open arrowheads and white arrowheads, respectively). Additionally, exocrine cells outside the pancreatic islets showed stronger PDI signals, and overlapping of NNAT and PDI signals was absent (Fig. 3C1–C5). Notably, endothelial cells with strong NNAT signals were negative for PDI (Fig. 3A, yellow arrow).

**Fig. 3.**
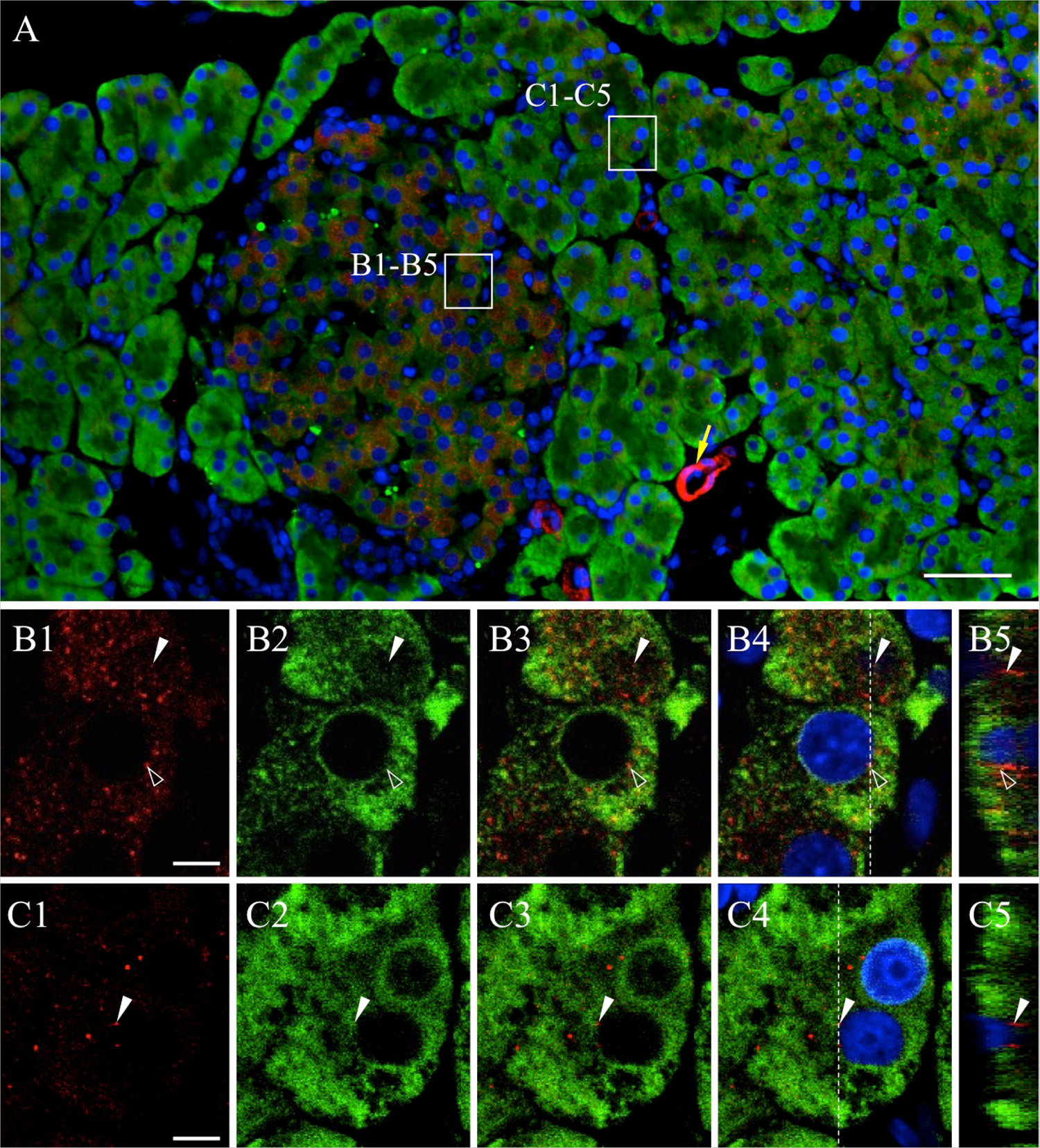
Intracellular localization of NNAT and protein disulfide isomerase (PDI) in the pancreas. Double immunostaining for NNAT (Cy3, red) and PDI (Cy5, green), an endoplasmic reticulum (ER) marker, in the rat pancreas on P60 was performed, and merged images with DAPI (blue) are shown. Boxed areas in a are enlarged in B1–B4 and C1–C4. Orthogonal projections by Z-stack imaging with 0.124-μm-thick slices are shown (B5, C5). White arrowhead: NNAT single-positive signal, open arrowhead: NNAT/PDI double-positive, yellow arrow: blood vessel. Scale bars = 50 μm (A), 5 μm (B, C).

In summary, NNAT signals were localized in some exocrine cells and in the majority of insulin-positive β-cells in the adult pancreas. In the middle stage of embryonic development, NNAT signals emerged in the area of exocrine cells, followed by in β-cells in the pancreatic islets and exocrine cells during late embryonic development. Analysis of subcellular localization revealed that NNAT was localized not only in the ER but also in other subcellular organelles.

### Immunohistochemistry for NNAT in the Tongue

The tongue and hair follicles are ectodermal appendages formed by interactions between epithelial cells derived from the surface ectoderm and the underlying mesenchyme. We next analyzed the anterior part of the tongue, which comprises two papillae, i.e., fungiform taste papilla housing taste buds and non-taste filiform papilla (Fig. 4, FFP and Flp, respectively). Immunohistochemistry for SOX2 in the tongue on E21.5 showed that the basal epithelial cells of both the filiform and fungiform papillae, and the taste bud cells in the fungiform papillae were positive for SOX2 (Fig. 4A1), showing weak and strong intensities of SOX2 signals, respectively (Fig. 4B1 and C1, double and single open arrowheads and white arrow, respectively), as previously reported.^21-23^ On P60, a number of SOX2-positive cells with elongated nuclei, characteristic of mature taste cells, were detected in the taste buds (Fig. 4D1, E1, yellow arrow). In addition, SOX2-positive progenitor cells with round nuclei were found in the basal compartment of the taste bud (Fig. 4C1, white arrow).

**Fig. 4.**
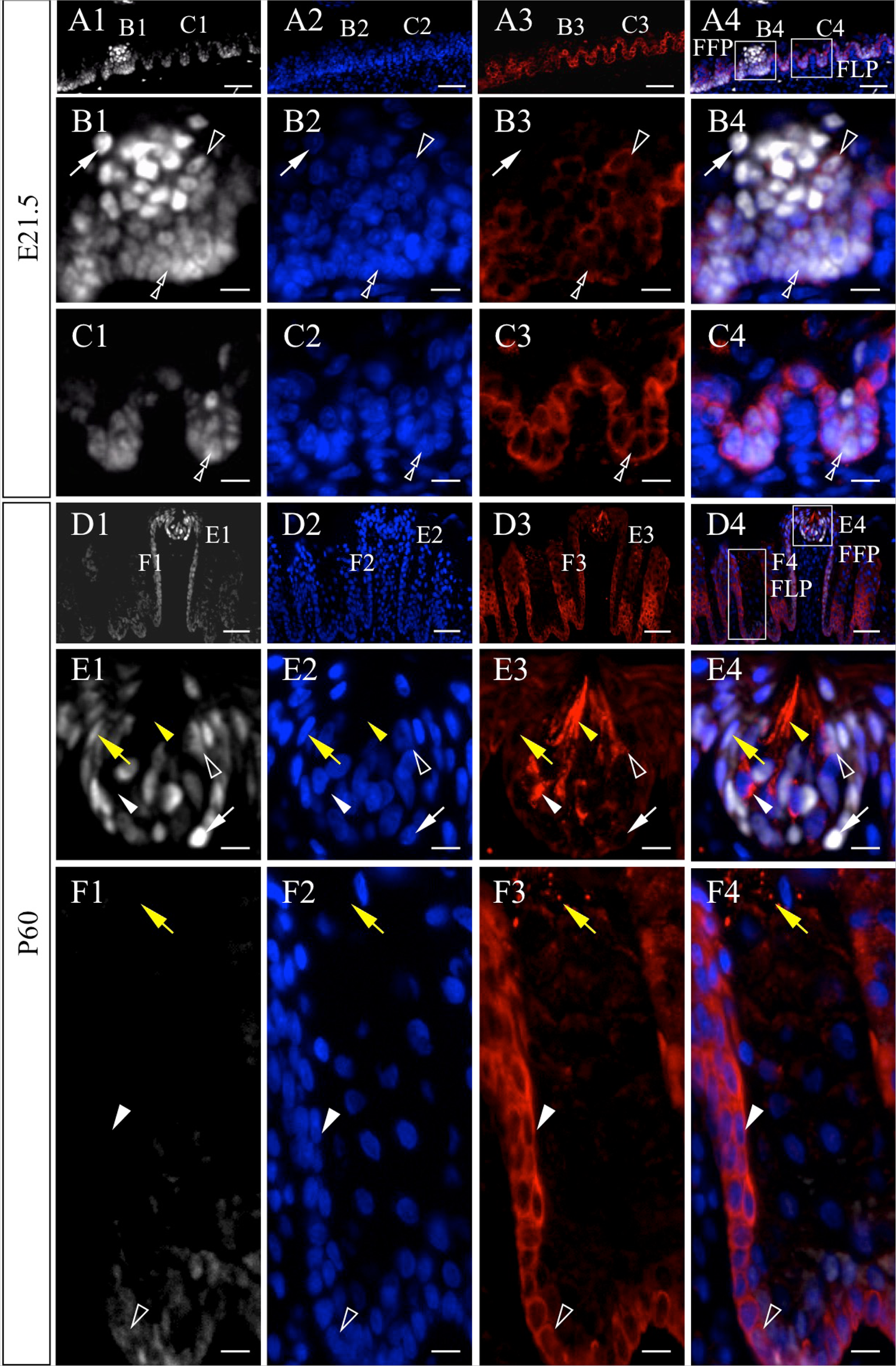
Immunohistochemistry for NNAT and SOX2 in embryonic and adult tongue tissues. Immunostaining for NNAT (red, Cy3) and SOX2 (white, Cy5) was performed for frozen sections from embryos on E21.5 (A1–A4) and P60 (D1–D4). Boxed areas in A4 and D4 are enlarged in B1–B4, C1–C4, E1–E4, and F1–F4. Nuclei are stained with DAPI (blue). White arrowhead: NNAT-single positive cell, open arrowhead: NNAT/SOX2- double positive cell, white arrow: SOX2-single positive cell, yellow arrowhead: NNAT- single positive elongated cell, yellow arrow: mature taste cell, double open arrowhead: basal epithelial cell. Flp: filiform papilla, FFP: fungiform papilla. Scale bars = 50 μm (A1–A4, D1–D4), 10 μm (B1–B4, C1–C4, E1–E4, F1–F4).

In the embryonic papillae on E21.5, double immunohistochemistry for NNAT and SOX2 showed that some NNAT/SOX2-double positive cells intermingled with SOX2-single positive cells in the developing taste buds (Fig. 4B1–B4, open arrowheads and white arrows, respectively). All SOX2-positive basal epithelial cells in the filiform papillae were positive for NNAT (Fig. 4C1–C4, double open arrowheads), as was also observed in the fungiform papillae (Fig. 4B1–B4, double open arrowheads).

In the adult fungiform papillae, various types of cell populations were present, i.e., NNAT-single-, NNAT/SOX2-double-, and SOX2-single-positive cells (Fig. 4E1–E4, white arrowheads, open arrowheads, and white arrows, respectively). Cells showing intense NNAT signals extended a characteristic elongated edge toward the taste pore (Fig. 4E1–E4, yellow arrowheads). In contrast, in the adult filiform papillae, although NNAT/SOX2-double positive cells were still observed in the basal epithelium at the bottom (Fig. 4F1–F4, open arrowheads), the upper portion of the basal epithelial layer was negative for SOX2 but positive for NNAT (Fig. 4F1–F4, white arrowheads). Notably, NNAT signals were present in the keratohyalin granules of cells existing in the interpapillary region of the filiform papillae (Fig. 4F1–F4, yellow arrows).

Double-immunostaining for NNAT and PDI was then performed to study the relationship between NNAT and the ER (Fig. 5A). Notably, cells in NNAT-positive domains in the filiform and fungiform papillae showed immature ER with weak PDI signals (Fig. 5B1–B4, D1–D4, open arrowheads), whereas intense PDI signals were observed in NNAT-negative keratinocytes in the filiform papilla (Fig. 5B1–B4, arrows) and in the fungiform papilla (Fig. D1–D4, arrows). NNAT-single positive anucleate cells were present in the corneocytes in the filiform papilla (Fig. 5C1-C4, white arrowheads).

**Fig. 5.**
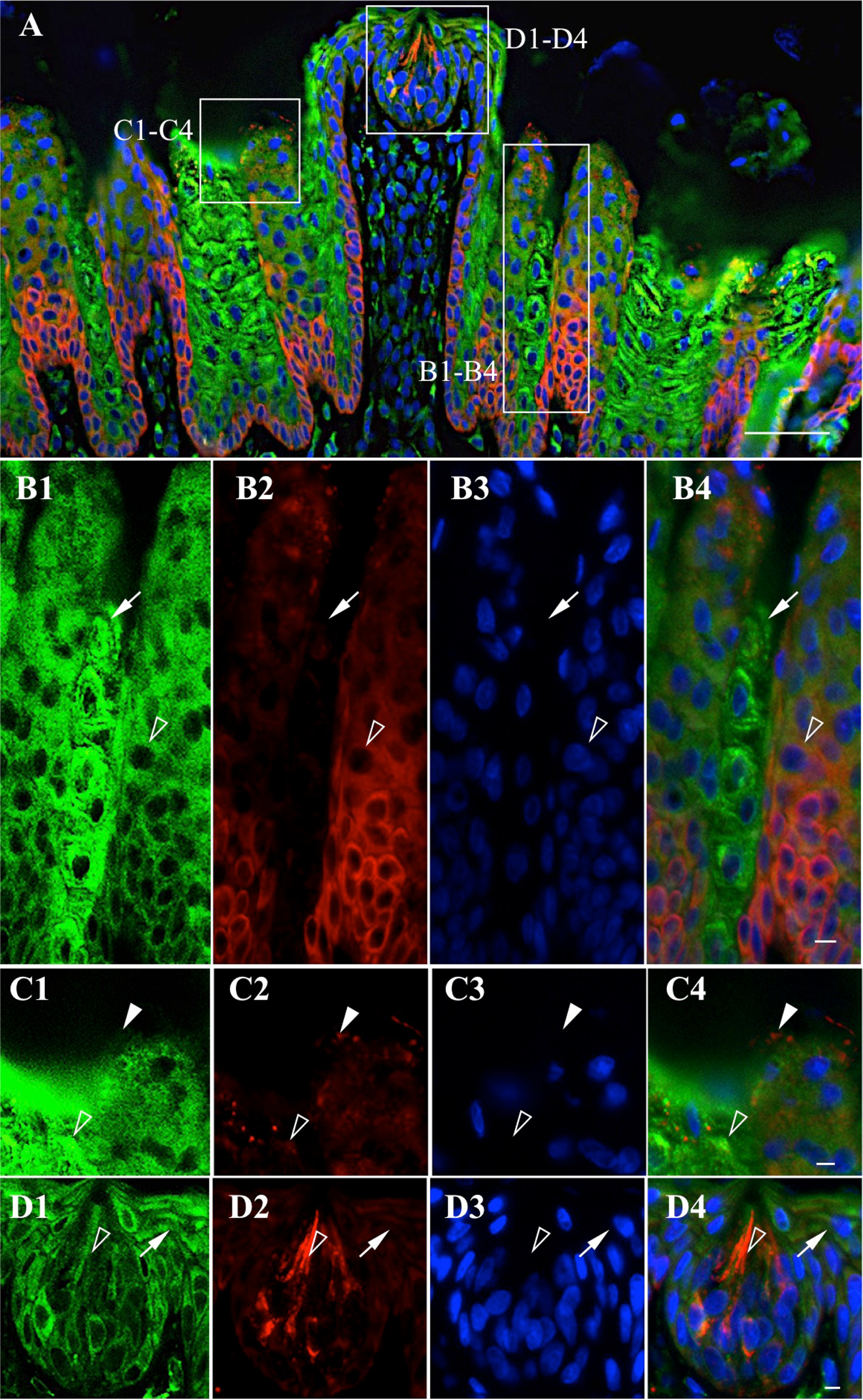
Intracellular localization of NNAT and PDI in filiform and fungiform papillae. Double immunostaining for NNAT and PDI in the rat filiform and fungiform papillae on P60 was performed. Boxed areas in a are enlarged in B1–B4, C1–C4, and D1–D4. NNAT and PDI were visualized with Cy3 (red) and Cy5 (green), respectively, and merged images with DAPI (blue) are shown. White arrowhead: NNAT-single positive signal, open arrowhead: NNAT/PDI-double positive signal, arrow: PDI-single positive signal. Scale bars = 50 μm (A), 5 μm (B4, C4, D4).

Thus, our findings showed that NNAT was localized in the basal epithelial cells and taste bud cells in the fungiform papillae and in both SOX2-positive and SOX2-negative basal epithelial cells in the filiform papillae. Cells in the NNAT-positive domain showed immature ER.

### Immunohistochemistry for NNAT in Whisker Hair Follicles

We next analyzed the whisker hair follicles of another ectodermal appendage. Hair follicle morphogenesis begins from the formation of the hair placode at the middle embryonic stage and continues until birth. Adult hair follicles are maintained by cycling through three phases of active hair follicle regeneration, i.e., hair growth (anagen), destruction (catagen), and rest (telogen). During full anagen phase, hair growth is active by differentiation of matrix progenitor cells of the hair bulb into the hair shaft, which is composed of three layers (i.e., the medulla, hair cortex, and hair cuticle), and the inner root sheath (IRS), which is composed of three layers (i.e., the IRS cuticle, Huxley’s layer, and Henle’s layer).

Immunostaining for NNAT was performed on E16.5 in rats, when the whisker hair follicles are still underdeveloped. All cells in the primordium were negative for NNAT (Fig. 6A, arrow). Longitudinal sections on E20.5 showed that positive signals for NNAT were present in the domain from the region close to the dermal papilla to the middle portion of the hair follicle (Fig. 6B). Cross-sections showed NNAT-positive concentric rings of IRS, composed of three layers (the IRS cuticle close to the hair shaft, the Huxley’s layer as middle layer, and the outermost Henle’s layer) essential for the shaping and guidance of the hair to the skin surface by keratinization (Fig. 6C, Ci, Hu, and He, respectively). The intensity of NNAT signals became stronger on P60 (Fig. 6D, E). Notably, the Henle’s layer in the outermost region and the cuticle layer in the innermost region of the IRS showed stronger NNAT signals than those in the middle of Huxley’s layer.

**Fig. 6.**
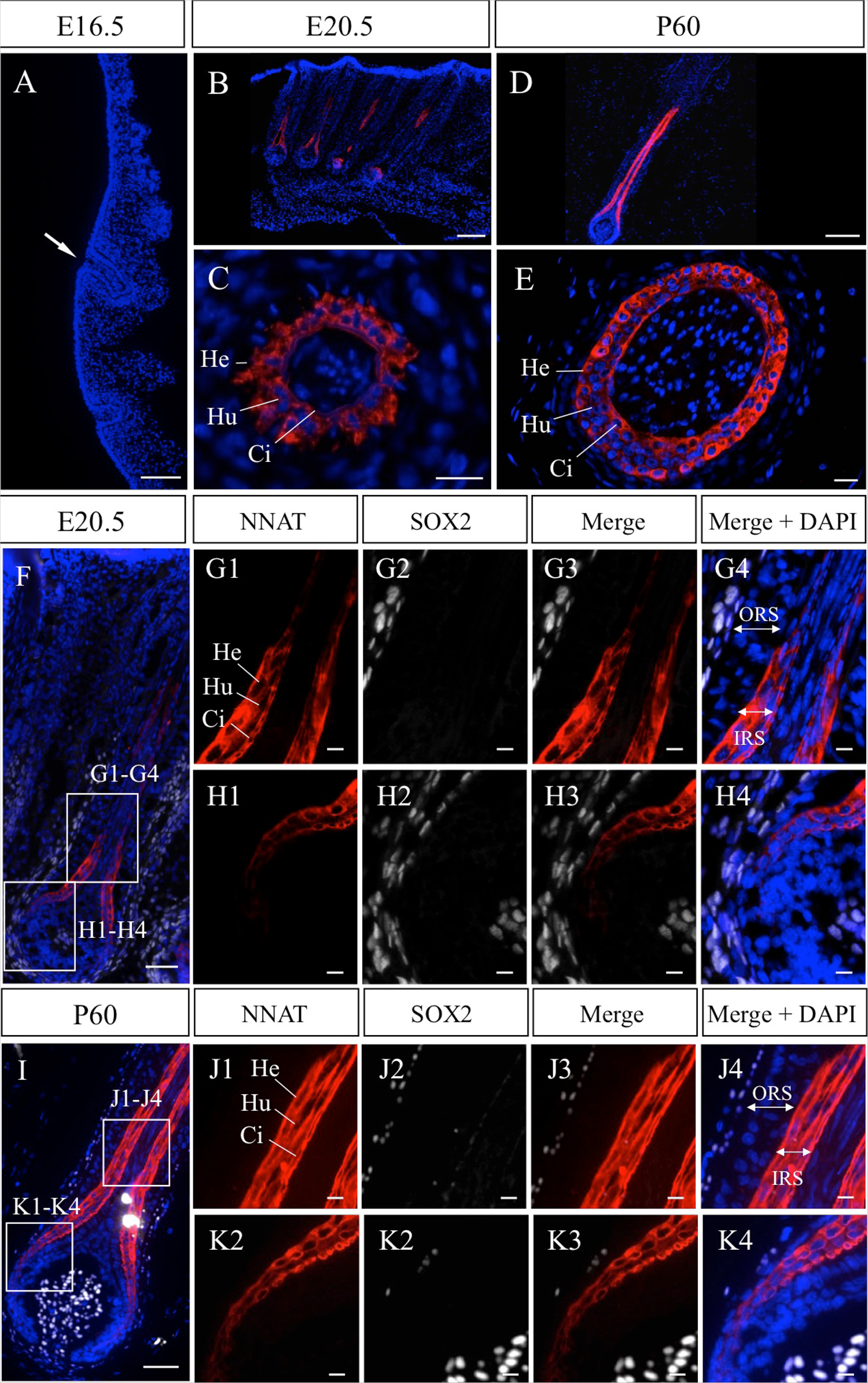
Immunohistochemistry for NNAT in embryonic and adult whisker hair follicles. Immunostaining for NNAT (red, Cy3) was performed using frozen sections prepared from embryos on E16.5 (A) and E20.5 (B, C) and from adult tissues on P60 (D, E). Longitudinal sections (A, B, D) and cross-sections (C, E) are shown. Nuclei are stained with DAPI (blue). Immunostaining for NNAT (red, Cy3) and SOX2 (white, Cy5) was performed for frozen sections prepared from tissues on E20.5 (F) and P60 (I). Boxed areas in F and I are enlarged in G1–G4, H1–H4, J1–J4, and K1–K4. Arrow indicates immature whisker hair follicle. He: Henle’s layer, Hu: Huxley’s layer, Ci: cuticle of inner root sheath, ORS: outer root sheath, IRS: inner root sheath. Scale bars = 50 μm (A–F, I), 10 μm (G1–G4, H1–H4, J1–J4, K1–K4).

Double-immunostaining for NNAT and SOX2 on E20.5 and P60 showed that SOX2-positive cells were localized in the dermal papilla (Fig. 6F, I). All NNAT-positive cells in the IRS and NNAT-negative cells in the outer root sheath (ORS) were negative for SOX2 (Fig. 6G1–G4, H1–H4, J1–J4, K1–K4).

Double-immunostaining for NNAT and PDI showed that PDI signals were widely observed in the cytoplasm of cells in the IRS (Fig. 7A). In addition to many signals overlapping with NNAT and PDI (Fig. 7B1–B4, C–C4, D–D4, E–E4, open arrowheads), NNAT signals negative for PDI and PDI signals negative for NNAT were also observed (Fig. 7B1–B4, C–C4, D–D4, E–E4, white arrowheads and arrows, respectively). Thus, these results demonstrated that NNAT-positive cells were limited to the IRS in the whisker hair follicles and that not all NNAT signals were located in the ER.

**Fig. 7.**
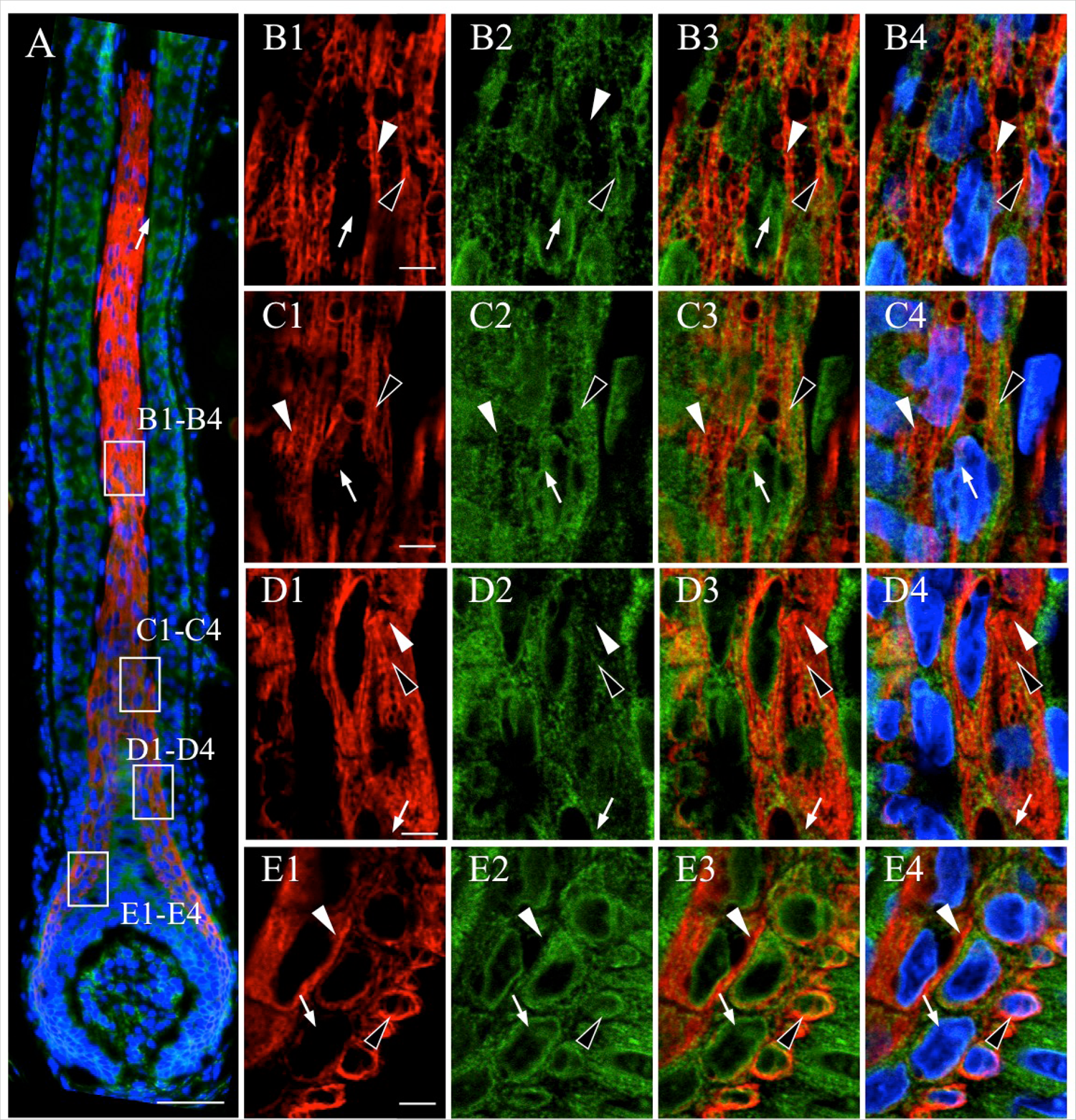
Intracellular localization of NNAT and PDI in whisker hair follicles. Double immunostaining for NNAT and PDI in rat whisker hair follicles on P60 was performed. Boxed areas in a are enlarged in B1–B4, C1–C4, D1–D4, and E1–E4. NNAT and PDI were visualized with Cy3 (red) and Cy5 (green), respectively, and merged images with DAPI (blue) are shown. White arrowhead: NNAT-single positive signal, open arrowhead: NNAT/PDI-double positive signal, arrow: PDI-single positive signal. Scale bars = 50 μm (A), 5 μm (B–E).

### Immunohistochemistry for NNAT in the Testis

The testes are reproductive tissues derived from the mesoderm and are composed of the seminiferous tubules and interstitium. The former is involved in spermatogenesis, in which spermatozoa are produced from spermatogonia, and the latter is involved in androgen production by Leydig cells.

In immature rat testes on P30, NNAT-positive cells were not observed (Fig. 8A1, A2). On P60, NNAT-positive cells appeared in the interstitium and seminiferous tubules of the testis (Fig. 8B). In the interstitium, granular NNAT signals were observed in Leydig cells, and the proportion of NNAT-positive cells was about 60% (Fig. 8C, D). In the seminiferous tubules, NNAT signals were absent from the spermatogonia, spermatocytes, and Sertoli cells (Fig. 8E1–M, SG, SP, and SC, respectively) during spermatogenesis. NNAT signals were first observed at acrosome vesicles in round spermatids after meiosis (Fig. 8K, RS) and showed aggregated localization across the crescent acrosome in elongated spermatids (Fig. 8M, ES). Even after spermatozoa moved to the epididymis, NNAT showed similar localization with slight attenuation of signal intensity between the caput epididymis and cauda epididymis (Fig. 8n and o, respectively).

**Fig. 8.**
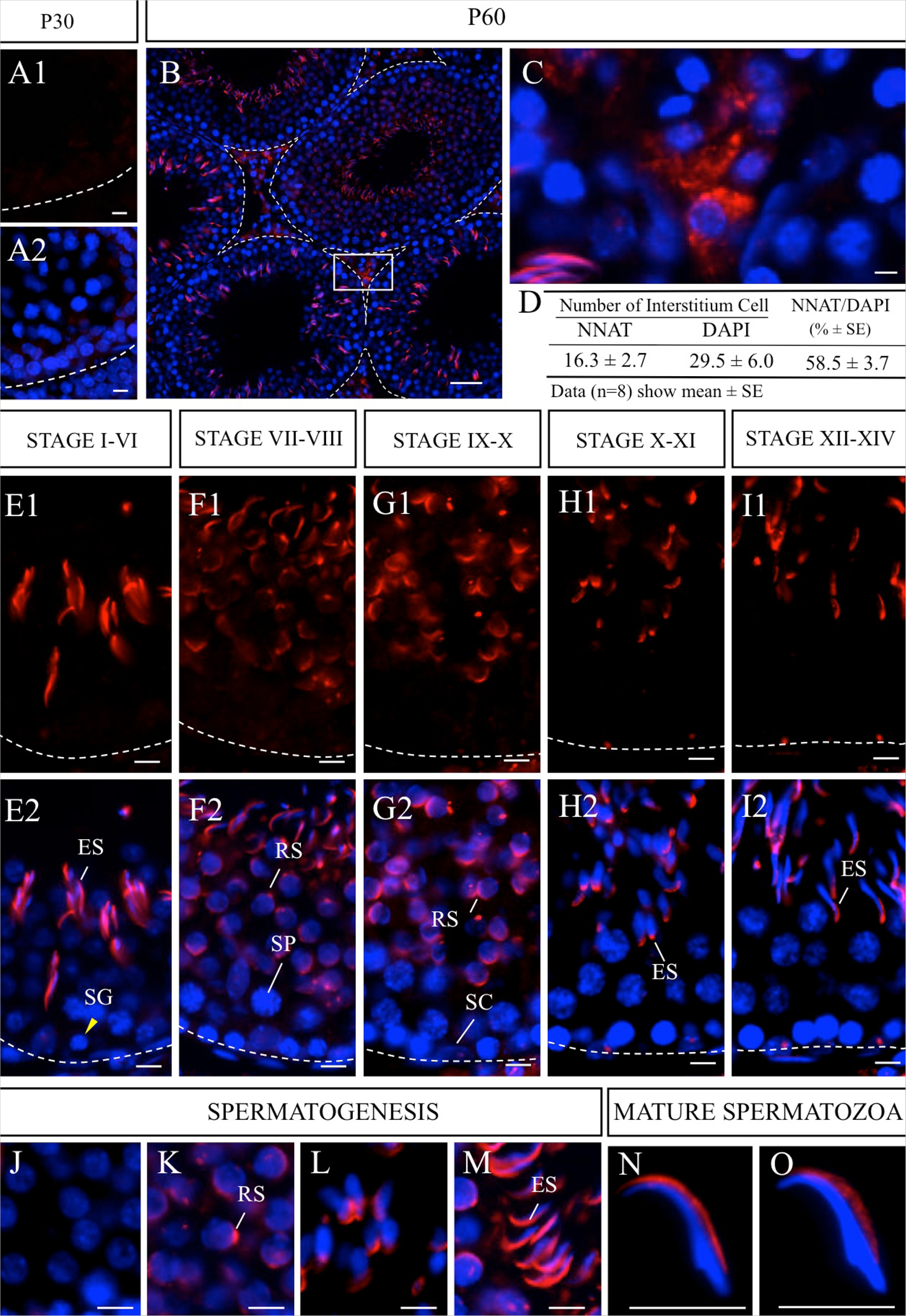
Immunohistochemistry for NNAT in immature and mature testes. Immunostaining for NNAT (red, Cy3) was performed for frozen sections prepared from immature (A1, A2, P30) and mature testes (B, C, P60). Developmental stages I–XIV in spermatogenesis (E1–J1) and merged images with DAPI (blue, E2–I2) are shown. Enlarged images of NNAT-positive cells during acrosome formation (J–M) and spermatozoa in the epididymal head (N) and cauda epididymis (O) are shown. Numbers and proportions of NNAT-positive cells in the testicular interstitium were counted for eight areas (19–36 cells each) and are shown in d. Data are presented as means ± SEs. ES: elongated spermatid, RS: round spermatid, SC: Sertoli cell, SG: spermatogonia, SP: spermatocyte. Scale bars = 50 μm (B), 5 μm (A1, A2, C, E, E2, F1, F2, G, G2, H1, H2,I1, I2, J–O).

Overall, these results confirmed that NNAT was specifically localized in the cytoplasm of Leydig cells and in the developing and mature acrosome. The correlation between NNAT and the ER was contradicted in the testes because spermatozoa are free from the ER and the acrosome is derived from the Golgi.^24^

## Discussion

NNAT is known to be involved in diverse intracellular roles in various tissues and/or cell types through Ca^2+^ signaling, glucose signaling, and inflammatory signaling.^25^ In contrast to its initial identification as an ER membrane protein,^8^ NNAT localization has also been observed in several subcellular organelles of the pituitary stem/progenitor cells^7^ and in the outer segment of rod photoreceptors, which do not contain an ER.^15^ The present immunohistochemical study demonstrated that NNAT was localized in developing tissues and in differentiating cells in the pancreas, tongue, whisker hair follicles, and testis and suggested that NNAT was involved in cell differentiation and in the function of terminally differentiated cells beyond the germ layer. In addition, our findings suggested that NNAT is not a true ER-associated protein.

In this study, we demonstrated, for the first time, that NNAT emerged in round spermatids, accompanied by acrosome formation, from the early stage in round spermatids to the maturation stage in spermatozoa. The ER is absent from spermatozoa, and the acrosome is derived from the Golgi apparatus.^24^ Moreover, in our previous study, we found that NNAT is localized in progenitor cells with rudimentary ER in the pituitary gland^7^ and the disk membrane of the outer segment of rod photoreceptors;^15^ thus, NNAT may be located in the membrane of appropriate subcellular organelles, not limited to the ER.

In whisker hair follicles, specific localization of NNAT in the IRS was shown. The IRS is known to abundantly produce trichohyalin,^26^ which mediates keratin filamentous assembly, although trichohyalin is also produced in the medulla of the hair shaft.^27^ In addition to trichohyalin in the IRS, several molecules involved in keratinization, such as peptidylarginine deiminase,^28^ involcurin,^29^ and transglutaminases,^30^ are produced. However, the localization of these molecules differs from that of NNAT, showing expression in other parts of the hair follicle in addition to the IRS. Notably, keratin K71, which has been reported to show limited localization in the three layers of the IRS,^31^ exhibits localization similar to that of NNAT. Deficiency of mK6irs1 protein, the mouse ortholog of keratin K71, causes alopecia.^32^ Thus, elucidation of the coexistence of NNAT and keratin K71 is essential in order to clarify the mechanisms mediating the development and keratinization of the IRS.

Sheridan et al.^33^ further described that the IRS is consistently negative for SERCA2, indicating that the Ca^2+^ concentration in the ER of the IRS is modulated by a mechanism that does not involve SERCA2. Accordingly, the interaction between NNAT and SERCA2 is cell- or tissue-specific, but not universal.

In the tongue, NNAT showed distinct localizations in undifferentiated and differentiated cells present in two types of papillae, i.e., filiform and fungiform papillae. In the filiform papillae, both SOX2-positive basal epithelial cells in the bottom region and SOX2-negative cells in the upper region were positive for NNAT, whereas filiform keratinocytes were negative for NNAT. These results indicated that NNAT-positive cells in the filiform papillae were stem/progenitor cells and committed cells to differentiate into keratinocytes. Moreover, NNAT was also localized in corneocytes, terminally differentiated keratinocytes. In contrast, in taste bud cells in the fungiform papillae, NNAT was localized in SOX2-positive basal epithelial cells and taste bud cells in addition to SOX2-negative mature taste cells. NNAT-positive taste cells negative for SOX2 showed extended NNAT signals toward the front edge of the elongated cells. The taste buds contain immature basal cells (type IV) and elongated mature taste cells with three morphologically distinguished types (types I, II, and III), which detect at least five types of tastes (salt, sweet, umami, bitter, and sour).^34^ Because type IV and type I cells express *Sox2* at high levels, whereas type II and type III cells rarely express *Sox2*,^23^ NNAT-positive taste receptor cells could be type II and/or III.

The molecular mechanisms of NNAT function are unclear. In type II cells in the taste buds, three tastes (sweet, umami, and bitter) are received through taste transduction mediated by taste receptors, TAS1Rs and TAS2Rs, which belong in the G protein-coupled receptor family. Notably, these receptors are frequently observed outside the gustatory system, such as in the sperm ^35^, and some are localized in acrosomes;^36^ in keratinocytes in the skin;^37^ airway, and gut;^38^ and in the placenta.^39^

In addition, one of the downstream signaling molecules of taste receptors, TRPM5, a Ca^2+^-activated cation channel, has been reported to localize in pancreatic β-cells.^40,41^ Therefore, to elucidate the role of NNAT, it may be necessary to clarify the signaling pathways involved in keratinization in the filiform papillae of the tongue and the IRS in the hair follicle, in acrosome formation in the spermatozoa, in photoreception in the rod photoreceptors, and in characteristic signals in NNAT-positive stem/progenitor cells.

In the pancreas, we showed that NNAT immunoreactive signals emerged in a few exocrine cells but not in insulin-positive cells in the middle stage of embryonic development, followed by signals in β-cells in the pancreatic islets in the late stage of embryonic development. In the postnatal pancreas, NNAT signals were localized in a few exocrine cells and the majority of β-cells. Millership et al.^42^ recently reported that the C-terminal hydrophilic domain of NNAT, located on the cytosolic side of the ER membrane, interacts with components of the signal peptidase complex to induce translocation of preproinsulin into the ER and to modulate insulin secretion in β-cells. Moreover, NNAT was shown to not be involved in Ca^2+^-mediated signaling using *Nnat*-null primary β-cells, suggesting that NNAT modulates insulin secretion at the stage of translocation of preproinsulin into the ER. However, the involvement of NNAT in secretion control may not be common to other endocrine tissues. In the pituitary gland, we observed that NNAT was present in stem/progenitor cells in the developing and developed pituitary but not in terminally differentiated hormone-producing cells.^7^ NNAT may have diverse roles in stem/progenitor cells and in terminally differentiated cells, even in such a relevant endocrine tissue. In addition, the roles of NNAT in insulin-negative cells of islet and exocrine cells and in subcellular organelles outside of the ER remain unclear.

In summary, our current findings showed that NNAT exhibited various localizations, suggesting diverse cell- or tissue-specific roles. NNAT functions beyond the germ layer and in the differentiating (tongue and testis) and terminally differentiated (pancreas, tongue, whisker hair follicle, and spermatozoa) cells. The observed subcellular localization of NNAT in the acrosome and inner root sheath provides important insights into the role of NNAT. Future studies are needed to identify molecules that interact with NNAT and explore the related molecular mechanisms.

## Acknowledgments

We wish to thank Dr. Hideji Yako and Dr. Masashi Higuchi for technical help and continuous support.

## Author Contributions

All authors have contributed to this article as follows: TK and YK developed the study design, coordination, and manuscript drafting; NK and SY performed immunohistochemical and quantitative analyses; TK and YK provided study supervision, as well as critical manuscript revision; and all authors have read and approved the manuscript as submitted.

## Literature Cited

1. Joseph R, Dou D, Tsang W. Neuronatin mRNA: alternatively spliced forms of a novel brain-specific mammalian developmental gene. Biomed Res.1995;690:92–98.

2. Aikawa S, Kato T, Elsaeesser F, Kato Y. Molecular cloning of porcine neuronatin and analysis of its expression during pituitary ontogeny. Exp Clin Endocrinol Diabetes.2003;111:475–79.

3. Joseph R, Dou D, Tsang W. Molecular cloning of a novel mRNA (neuronatin) that is highly expressed in neonatal mammalian brain. Biochem Biophys Res Commun.1994;201:1227–34.

4. Usui H, Ichikawa T, Miyazaki Y, Nagai S, Kumanishi T. Isolation of cDNA clones of the rat mRNAs expressed preferentially in the prenatal stages of brain development. Brain Res Dev Brain Res.1996;97:185–93.

5. Wijnholds J, Chowdhury K, Wehr R, Gruss P. Segment-specific expression of the neuronatin gene during early hindbrain development. Dev Biol.1995;171:73–84.

6. Nishida Y, Yoshioka M, St-Amand J. The top 10 most abundant transcripts are sufficient to characterize the organs functional specificity: evidences from the cortex, hypothalamus and pituitary gland. Gene.2005;344:133–41.

7. Kanno N, Higuchi M, Yoshida S, Yako H, Chen M, Ueharu H, Nishimura N, Mitsuishi H, Kato T, Kato Y. Expression studies of Neuronatin in the prenatal and postnatal rat pituitary. Cell Tissue Res.2016;364:273–88.

8. Joe MK, Lee HJ, Suh YH, Han KL, Lim JH, Song J, Seong JK, Jung MH. Crucial roles of neuronatin in insulin secretion and high glucose-induced apoptosis in pancreatic beta-cells. Cell Signal.2008;20:907–15.

9. Lin HH, Bell E, Uwanogho D, Perfect LW, Noristani H, Bates TJ, Snetkov V, Price J, Sun YM. Neuronatin promotes neural lineage in ESCs via Ca(2+) signaling. Stem Cells.2010;28:1950–60.

10. Dugu L, Nakahara T, Wu Z, Uchi H, Liu M, Hirano K, Yokomizo T, Furue M. Neuronatin is related to keratinocyte differentiation by up-regulating involucrin. J Dermatol Sci.2014;73:225–31.

11. Suh YH, Kim WH, Moon C, Hong YH, Eun SY, Lim JH, Choi JS, Song J, Jung MH. Ectopic expression of Neuronatin potentiates adipogenesis through enhanced phosphorylation of cAMP-response element-binding protein in 3T3-L1 cells. Biochem Biophys Res Commun.2005;337:481–89.

12. Oyang EL, Davidson BC, Lee W, Poon MM. Functional characterization of the dendritically localized mRNA neuronatin in hippocampal neurons. PLoS One.2011;6:e24879.

13. Ryu S, McDonnell K, Choi H, Gao D, Hahn M, Joshi N, Park SM, Catena R, Do Y, Brazin J, Vahdat LT, Silver RB, Mittal V. Suppression of miRNA-708 by polycomb group promotes metastases by calcium-induced cell migration. Cancer Cell.2013;23:63–76.

14. Yang J, Wei J, Wu Y, Wang Z, Guo Y, Lee P, Li X. Metformin induces ER stress-dependent apoptosis through miR-708-5p/NNAT pathway in prostate cancer. Oncogenesis.2015;4:e158.

15. Shinde V, Pitale PM, Howse W, Gorbatyuk O, Gorbatyuk M. Neuronatin is a stress-responsive protein of rod photoreceptors. Neuroscience.2016;328:1–8.

16. Gburcik V, Cleasby ME, Timmons JA. Loss of neuronatin promotes “browning” of primary mouse adipocytes while reducing Glut1-mediated glucose disposal. Am J Physiol Endocrinol Metab.2013;304:E885–94.

17. Sharma J, Rao SN, Shankar SK, Satishchandra P, Jana NR. Lafora disease ubiquitin ligase malin promotes proteasomal degradation of neuronatin and regulates glycogen synthesis. Neurobiol Dis.2011;44:133–41.

18. Sharma J, Mukherjee D, Rao SN, Iyengar S, Shankar SK, Satishchandra P, Jana NR. Neuronatin-mediated aberrant calcium signaling and endoplasmic reticulum stress underlie neuropathology in Lafora disease. J Biol Chem.2013;288:9482–90.

19. Chu K, Tsai M-J. Neuronatin, a downstream target of BETA2/NeuroD1 in the pancreas, is involved in glucose-mediated insulin secretion. Diabetes.2005;54:1064–73.

20. Mzhavia N, Yu S, Ikeda S, Chu TT, Goldberg I, Dansky HM. Neuronatin: a new inflammation gene expressed on the aortic endothelium of diabetic mice. Diabetes.2008;57:2774–83.

21. Okubo T, Clark C, Hogan BL. Cell lineage mapping of taste bud cells and keratinocytes in the mouse tongue and soft palate. Stem Cells.2009;27:442–50.

22. Okubo T, Pevny LH, Hogan BL. Sox2 is required for development of taste bud sensory cells. Genes Dev.2006;20:2654–59.

23. Suzuki Y. Expression of Sox2 in mouse taste buds and its relation to innervation. Cell Tissue Res.2008;332:393–401.

24. Nakamura M, Moriya M, Baba T, Michikawa Y, Yamanobe T, Arai K, Okinaga S, Kobayashi T. An endoplasmic reticulum protein, calreticulin, is transported into the acrosome of rat sperm. Exp Cell Res.1993;205:101–10.

25. Pitale PM, Howse W, Gorbatyuk M. Neuronatin Protein in Health and Disease. J Cell Physiol.2017;232:477–81.

26. Birbeck MS, Mercer EH. The electron microscopy of the human hair follicle. III. The inner root sheath and trichohyaline. J Biophys Biochem Cytol.1957;3:223–30.

27. Hamilton EH, Payne RE, Jr., O’Keefe EJ. Trichohyalin: presence in the granular layer and stratum corneum of normal human epidermis. J Invest Dermatol.1991;96:666–72.

28. Rogers G, Winter B, McLaughlan C, Powell B, Nesci T. Peptidylarginine deiminase of the hair follicle: characterization, localization, and function in keratinizing tissues. J Invest Dermatol.1997;108:700–7.

29. Thibaut S, Candi E, Pietroni V, Melino G, Schmidt R, Bernard BA. Transglutaminase 5 expression in human hair follicle. J Invest Dermatol.2005;125:581–5.

30. Alibardi L. Ultrastructural immunolocalization of involucrin in the medulla and inner root sheath of the human hair. Ann Anat.2012;194:345–50.

31. Langbein L, Yoshida H, Praetzel-Wunder S, Parry DA, Schweizer J. The keratins of the human beard hair medulla:the riddle in the middle. J Invest Dermatol.2010;130:55–73.

32. Peters T, Sedlmeier R, Bussow H, Runkel F, Luers GH, Korthaus D, Fuchs H, Hrabe de Angelis M, Stumm G, Russ AP, Porter RM, Augustin M, Franz T. Alopecia in a novel mouse model RCO3 is caused by mK6irs1 deficiency. J Invest Dermatol.2003; 121:674–80.

33. Sheridan AT, Hollowood K, Sakuntabhai A, Dean D, Hovnanian A, Burge S. Expression of sarco/endo-plasmic reticulum Ca2+-ATPase type 2 isoforms (SERCA2) in normal human skin and mucosa, and Darier’s disease skin. Br J Dermatol.2002;147:670–4.

34. Sullivan JM, Borecki AA, Oleskevich S. Stem and progenitor cell compartments within adult mouse taste buds. Eur J Neurosci.2010;31:1549–60.

35. Li F. Taste perception:from the tongue to the testis. Mol Hum Reprod.2013;19:349–60.

36. Meyer D, Voigt A, Widmayer P, Borth H, Huebner S, Breit A, Marschall S, de Angelis MH, Boehm U, Meyerhof W, Gudermann T, Boekhoff I. Expression of Tas1 taste receptors in mammalian spermatozoa: functional role of Tas1r1 in regulating basal Ca(2)(+) and cAMP concentrations in spermatozoa. PLoS One.2012;7:e32354.

37. Wolfle U, Elsholz FA, Kersten A, Haarhaus B, Muller WE, Schempp CM. Expression and functional activity of the bitter taste receptors TAS2R1 and TAS2R38 in human keratinocytes. Skin Pharmacol Physiol.2015;28:137–46.

38. Finger TE, Kinnamon SC. Taste isn’t just for taste buds anymore. F1000 Biol Rep.2011;3:20.

39. Wolfle U, Elsholz FA, Kersten A, Haarhaus B, Schumacher U, Schempp CM. Expression and Functional Activity of the Human Bitter Taste Receptor TAS2R38 in Human Placental Tissues and JEG-3 Cells. Molecules.2016;21:306.

40. Prawitt D, Monteilh-Zoller MK, Brixel L, Spangenberg C, Zabel B, Fleig A, Penner R. TRPM5 is a transient Ca2+-activated cation channel responding to rapid changes in [Ca2+]i. Proc Natl Acad Sci U S A.2003;100:15166–71.

41. Philippaert K, Pironet A, Mesuere M, Sones W, Vermeiren L, Kerselaers S, Pinto S, Segal A, Antoine N, Gysemans C, Laureys J, Lemaire K, Gilon P, Cuypers E, Tytgat J, Mathieu C, Schuit F, Rorsman P, Talavera K, Voets T, Vennekens R. Steviol glycosides enhance pancreatic beta-cell function and taste sensation by potentiation of TRPM5 channel activity. Nat Commun.2017;8:14733.

42. Millership SJ, da Silva Xavier G, Choudhury AI, Bertazzo S, Chabosseau P, Pedroni SM, Irvine EE, Montoya A, Faull P, Taylor WR, Kerr-Conte J, Pattou F, Ferrer J, Christian M, John RM, Latreille M, Liu M, Rutter GA, Scott J, Withers DJ. Neuronatin regulates pancreatic beta cell insulin content and secretion. J Clin Invest.2018;128:3369–81.

